# Estimating time of HIV-1 infection from next-generation sequence diversity

**DOI:** 10.1101/129387

**Authors:** Vadim Puller, Richard Neher, Jan Albert

**Affiliations:** Max Planck Institute for Developmental Biology, 72076 Tübingen, Germany; Biozentrum, University of Basel, Basel, Switzerland; SIB Swiss Institute of Bioinformatics, Basel, Switzerland; Department of Microbiology, Tumor and Cell Biology, Karolinska Institute, Stockholm, Sweden; Department of Clinical Microbiology, Karolinska University Hospital, Stockholm, Sweden

**Author notes:** Current Address: Biozentrum, University of Basel, Basel, Switzerland (Vadim Puller, Richard Neher).

## Abstract

Estimating the time since infection (TI) in newly diagnosed HIV-1 patients is challenging, but important to understand the epidemiology of the infection. Here we explore the utility of virus diversity estimated by next-generation sequencing (NGS) as novel biomarker by using a recent genome-wide longitudinal dataset obtained from 11 untreated HIV-1-infected patients with known dates of infection. The results were validated on a second dataset from 31 patients.

Virus diversity increased linearly with time, particularly at 3rd codon positions, with little inter-patient variation. The precision of the TI estimate improved with increasing sequencing depth, showing that diversity in NGS data yields superior estimates to the number of ambiguous sites in Sanger sequences, which is one of the alternative biomarkers. The full advantage of deep NGS was utilized with continuous diversity measures such as average pairwise distance or site entropy, rather than the fraction of polymorphic sites. The precision depended on the genomic region and codon position and was highest when 3rd codon positions in the entire *pol* gene were used. For these data, TI estimates had a mean absolute error of around 1 year. The error increased only slightly from around 0.6 years at a TI of 6 months to around 1.1 years at 6 years.

Our results show that virus diversity determined by NGS can be used to estimate time since HIV-1 infection many years after the infection, in contrast to most alternative biomarkers. We provide the regression coefficients as well as web tool for TI estimation.

**Author summary:** HIV-1 establishes a chronic infection, which may last for many years before the infected person is diagnosed. The resulting uncertainty in the date of infection leads to difficulties in estimating the number of infected but undiagnosed persons as well as the number of new infections, which is necessary for developing appropriate public health policies and interventions. Such estimates would be much easier if the time since HIV-1 infection for newly diagnosed cases could be accurately estimated. Three types of biomarkers have been shown to contain information about the time since HIV-1 infection, but unfortunately, they only distinguish between recent and long-term infections (concentration of HIV-1-specific antibodies) or are imprecise (immune status as measured by levels of CD4+ T-lymphocytes and viral sequence diversity measured by polymorphisms in Sanger sequences).

In this paper, we show that recent advances in sequencing technologies, i.e. the development of next generation sequencing, enable significantly more precise determination of the time since HIV-1 infection, even many years after the infection event. This is a significant advance which could translate into more effective HIV-1 prevention.

## Introduction

At diagnosis, most HIV-1 infected patients have an established HIV-1 infection of unknown duration. This uncertainty complicates inference about the epidemiology of HIV-1. Consequently, there is limited information about the true incidence of HIV-1, the number of hidden, undiagnosed infected persons, the magnitude of the problem referred to as “late presentation” and other important aspects of HIV-1 spread.

Several biomarkers that classify patients as recently or long-term infected have been used to estimate HIV-1 incidence in populations [1–7]. These biomarkers can be divided into three main categories: (i) serological incidence tests, (ii) CD4+ T-lymphocyte (CD4)-based estimates and (iii) sequence-based estimates. Importantly, these biomarkers usually do not determine the time since infection (TI), which limits their utility.

Serological incidence assays are based on knowledge about the development and maturation of HIV-1 antibody responses (reviewed in [1, 4–6, 8]). Among the serological assays, the BED assay and the LAg avidity assay are the most widely used [4, 8–10]. CD4 counts are determined as part of routine clinical care, a CD4 count below 350 cells/*μ*L (or an AIDS-defining illness) at diagnosis is defined as late presentation [11, 12]. However, CD4 count is an imprecise measure of TI because its rate of decline is quite variable [13–16].

Sequence-based methods focus on the increase in intra-patient HIV-1 sequence diversity following infection [17]. Kouyos et al [18] showed that time since infection correlated with the fraction of polymorphic nucleotides in partial HIV-1 *pol* gene sequences determined by Sanger sequencing. Others have later reported similar findings [19–21]. This idea was expanded to other measures of sequence diversity, such as mean Hamming distance [6, 7, 22] and high-resolution melting (HRM) [23]. These studies (except HRM) used sequences generated by traditional Sanger population sequencing often performed as part of routine HIV-1 resistance testing.

Here, we have investigated the utility of estimating time since HIV-1 infection using genetic diversity in whole genome deep sequencing data generated by next-generation sequencing (NGS) on the Illumina platform [24]. We show that sequence diversity is a useful biomarker that grows approximately linearly with time during the first 8 years of infection. We found that the *pol* gene was best suited to calculate TI because diversity, mostly at third positions, accumulated more steadily in *pol* than in other genomic regions. Inclusion of intra-patient single nucleotide variants (iSNVs, also referred to as “polymorphisms”) down to the detection limit of NGS (i.e. 0.3%) improved the accuracy of TI estimations as did exclusion of 1st and 2nd codon position sites. NGS provided more accurate estimates of TI than Sanger sequencing, which at best detects iSNVs down to 25% [25–27].

## Materials and methods

### Patients

The study included two sets of HIV-1 whole-genome sequence data; a training dataset and a validation dataset.

The training dataset consisted of sequence data from 11 patients who were diagnosed in Sweden between 1990 and 2003 (Table 1 and S6 Table). Details about the patients and the samples have been published [24] and are also available online at hiv.biozentrum.unibas.ch. The patients were selected based on the following criteria: 1) A relatively well-defined time of infection based on a laboratory-confirmed primary HIV-1 infection (PHI) or a negative HIV antibody test less than two years before the first positive test; 2) No antiretroviral therapy (ART) during a minimum of approximately five years following diagnosis; and 3) Availability of biobank plasma samples covering this time period. As previously described 6 - 12 plasma samples per patient were retrieved from biobanks and used for full-genome HIV-1 RNA deep sequencing [24].

**Table 1.**
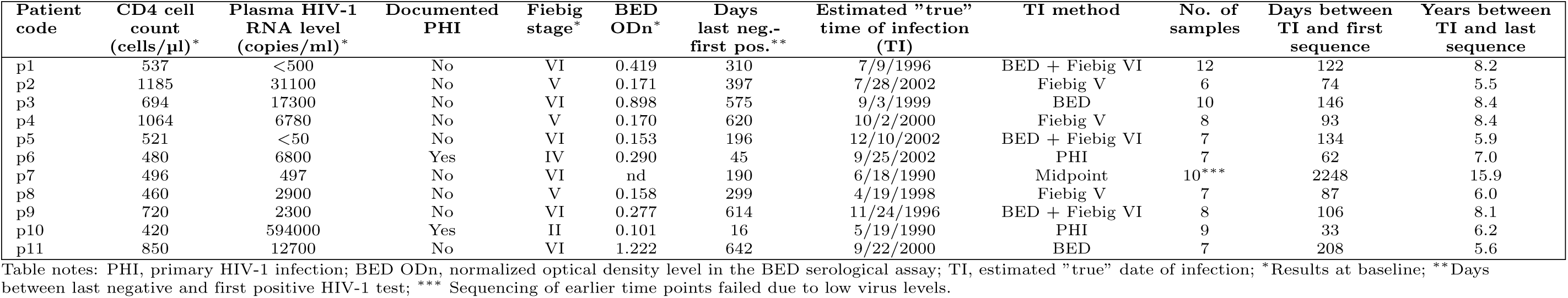
Summary of patient characteristics.

The validation dataset consisted of data from 31 patients who were diagnosed in Sweden between 2003 and 2010 (S7 Table). The patients were selected using similar criteria as the 11 patients in the training dataset, but the follow-up time without ART was shorter (median 2.9 years, range 1.4-6.2 years). For each patient, one plasma sample collected early during follow-up and one plasma sample collected late during follow-up was deep sequenced using the same method as for the 11 patients.

### Ethics statement

The study was conducted according to the Declaration of Helsinki. Ethical approval was granted by the Regional Ethical Review Board in Stockholm, Sweden (registration no. 2012-505 and 2014-646 for the validation dataset and 2002-367, 2004-797, 2007-1533 and 2011-1854 for the training dataset). Patients participating in the study gave written and oral informed consent to participate.

### Determination of the “true” time of infection (TI)

For each patient in the training and validation datasets the “true” time of infection (TI) was estimated using a hierarchical scheme based on clinical and laboratory findings as previously described [24, 28]:

1. Laboratory-confirmed PHI. Infection was considered to have occurred 17 days prior to date of first hospital visit based on an average incubation time from infection to development of PHI of 14 days [29] and an estimated patient delay of 3 days.
2. Fiebig staging [28] was used if the necessary laboratory results were available and the patient found to be in Fiebig stage I–V, which were considered to correspond to 13, 18, 22, 27 and 65 days since infection based on Cohen et al. [30].
3. BED assay results (i.e. normalized optical density, ODn) analyzed with a published time-continuous model of development of BED-reactive antibodies [31] if the ODn level corresponded to *<*1 year since infection after which ODn levels start to saturate.
4. Midpoint between the last negative and the first positive HIV test.

If possible information from several methods to determine TI were combined. The true TIs were considered to be without measurement error. Comprehensive information is provided in the S6 Table and the S7 Table.

### CD4 counts, virus levels and BED tests

Plasma HIV-1 RNA levels were measured using the Cobas AmpliPrep sample preparation system followed by analysis using the Cobas Amplicor HIV-1 monitor version 1.5 or the Cobas TaqMan HIV-1 v1.0 or v2.0 (Roche Molecular Systems, Basel, Switzerland). CD4+ T-lymphocyte (CD4) cells were enumerated using flow cytometry.

As part of determination of the true TIs, BED testing was performed on the first plasma sample from each study subject using the Aware BED EIA HIV-1 Incidence Test (Calypte Biomedical Corporation, Portland, OR, USA) according to the manufacturer’s instructions.

### HIV-1 RNA sequences

Whole-genome deep-sequencing of virus RNA populations in plasma samples obtained before start of therapy was performed as previously described [24, 32]. In short, total RNA in plasma was extracted using RNeasy Lipid Tissue Mini Kit (Qiagen Cat No. 74804) and amplified using a one-step RT-PCR with outer primers for six overlapping regions and Superscript III One-Step RT-PCR with Platinum Taq High Fidelity High Enzyme Mix (Invitrogen, Carlsbad, California, US). An optimized Illumina Nextera XT library preparation protocol was used together with a kit from the same supplier to build DNA libraries, which were sequenced on the Illumina MiSeq instrument with 2x250bp or 2x300bp sequencing kits (MS-102-2003/MS-10-3003).

Sequencing reads are available in the European Nucleotide Archive under accession number PRJEB9618 and processed data are available at hiv.biozentrum.unibas.ch.

The samples of the validation dataset were sequenced and processed using the same protocol and analysis pipeline [24, 32]. Patient-specific consensus sequences were constructed using an iterative mapping procedure. All reads were then remapped against this reference to calculate iSNV frequencies (i.e. pile-ups or tables how often each nucleotide was observed at every position of the genome). Sequencing and mapping/assembly was successful for 56 of the 62 samples. Filtered short-reads were submitted to ENA and are available under study accession number PRJEB21629. iSNV frequency counts at each position of *pol* and *gag* are available as part of the analysis code repository at github.com/neherlab/HIV_time_of_infection.

All analyses were done in Python using the libraries numpy, biopython, and matplotlib [33–35]. These iSNV frequency tables were then used to calculate average pairwise distances, average alignment entropies, or the number of sites with variation above a cutoff *x*_*c*_.

### Statistical procedures

We have used three different diversity measures: fraction of polymorphic sites, average pairwise distance per length, and site entropy. All of these measures can be straight-forwardly calculated from the frequencies of different nucleotides *x*_*iα*_ at site *i* = 1 *… L* and *α* ∊ {A, C, G, T} along the genome. Prior to calculation the nucleotide frequencies for each site were normalized to sum to unity (i.e. ignoring gaps or positions not called by the sequencer.)

For all methods, we introduce a cutoff *x*_*c*_. Sites at which the sum of all minor variants is smaller than *x*_*c*_ contribute zero to the diversity measure. This cutoff serves to filter sequencing errors or rare variation that cannot be reproducibly measured across samples. When using the *fraction of polymorphic sites* as diversity measure, *x*_*c*_ serves as the value above which sites are considered “polymorphic”. Specifically, the *fraction of polymorphic sites* is defined as

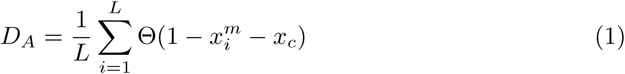

where 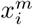 is the frequency of the dominant nucleotide at position *i*, and ⊝(*x*) is 1 when *x >* 0 and 0 otherwise (i.e. 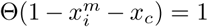 when 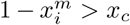 and 0 otherwise). *D*_*A*_ is thus the fraction of sites at which the dominant nucleotide is less frequent than 1 − *x*_*c*_.

The *average pairwise distance* is the probability that two randomly drawn sequences have different nucleotides at a specified position, averaged over all positions. It can be calculated from the *x*_*iα*_ as

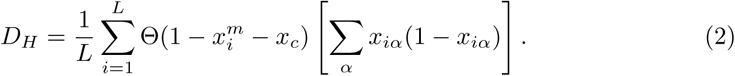

The quantity defined by Eq. (2) is the conventional Nei-Li nucleotide diversity [36] (∑_*α*_ *x*_*iα*_ (1 − *x*_*iα*_)) averaged over the sites. We refer to it as “average pairwise distance” whenever it is necessary to distinguish it from the other diversity measures introduced here, but otherwise call it simply “diversity”.

This diversity measure is similar to the fraction of polymorphic sites with the important difference that the contribution of each site is weighted by a frequency dependent factor.

The *average site entropy* is defined by

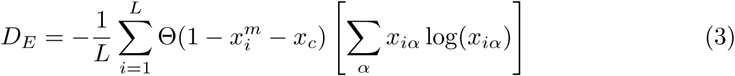

and differs from the average pairwise distance by the weighing function used. The entropy puts more weight on sites with rare variation. This can increase the information of the measure about TI, but can also be detrimental if too much weight is put on frequencies dominated by sequencing error. We evaluate and discuss the merits of the different measures below. We use the average pairwise distance as default diversity measure.

Given a diversity measure *D*, we model TI by

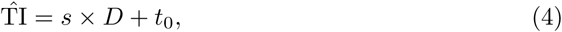

where *s* is the conversion factor between diversity and time, and *t*_0_ is the intercept value intended to accommodate possible non-linearity of diversity at small times.

To estimate values of *s, t*_0_ we minimized the average prediction error for the available data points in respect to these two parameters (see S1 Appendix for more details) The error in estimating *s, t*_0_ was calculated by randomly sampling the patients (bootstrapping over the patients).

### Cross-validation

To test the accuracy of the TI inference we used ten of the eleven patients as training data (to determine the slope and the intercept) treating the eleventh patient as test data. This procedure was repeated for every patient. Leaving out one patient at a time, rather than one sample at a time, gives more accurate confidence intervals as different samples from the same patients are correlated. We included in statistical analysis only samples where more than 50% of the sites in the averaging window were successfully sequenced to minimal coverage of 100.

Additional validation was provided by applying the slopes and intercepts obtained by analyzing the data from the training set of 11 patients to the validation dataset of 31 other patients with known times of infection.

## Results

### Patients characteristics

The training dataset consisted of recently published longitudinal full-genome deep sequencing data from 11 HIV-1 infected patients who were diagnosed in Sweden between 1990 and 2003 (6-12 samples per patient) and had a relatively well-defined time of infection (TI) [24]. The patient characteristics are summarized in Table 1 and fully described in the S6 Table. Nine of the eleven patients were MSM infected with HIV-1 of genetic subtype B. TI was hierarchically estimated using clinical and laboratory data (see Methods). Here, we take this estimate as the true time of infection and investigate how accurately TI can be estimated from sequence diversity in one sample. We will refer to this estimate as the *estimated time since infection* (ETI).

To validate the findings we used a second dataset consisting of similar sequence data from 31 additional patients (two samples per patient). The patient characteristics in validation dataset was more diverse than for the 11 patients training dataset. Thus, 16 of the 31 patients were infected with non-B-subtypes of HIV-1 and 11 patients belonged to other transmission groups than MSM. See methods and (S7 Table).

### Sequence diversity as a biomarker

All three diversity measures described in Materials and Methods grew linearly with time in the eleven patients. Fig. 1 shows average pairwise distance in the *pol* separately for each codon position. Most diversity in *pol* is synonymous and accumulates at 3rd codon positions, while diversity at 1st and 2nd codon positions remained low throughout. This pattern was less pronounced in other genes [24, 37] (see also S1 Fig and S2 Fig). In *env*, in particular, frequent selective sweeps result in a saturation of diversity later in infection [17, 24].

**Fig 1.**
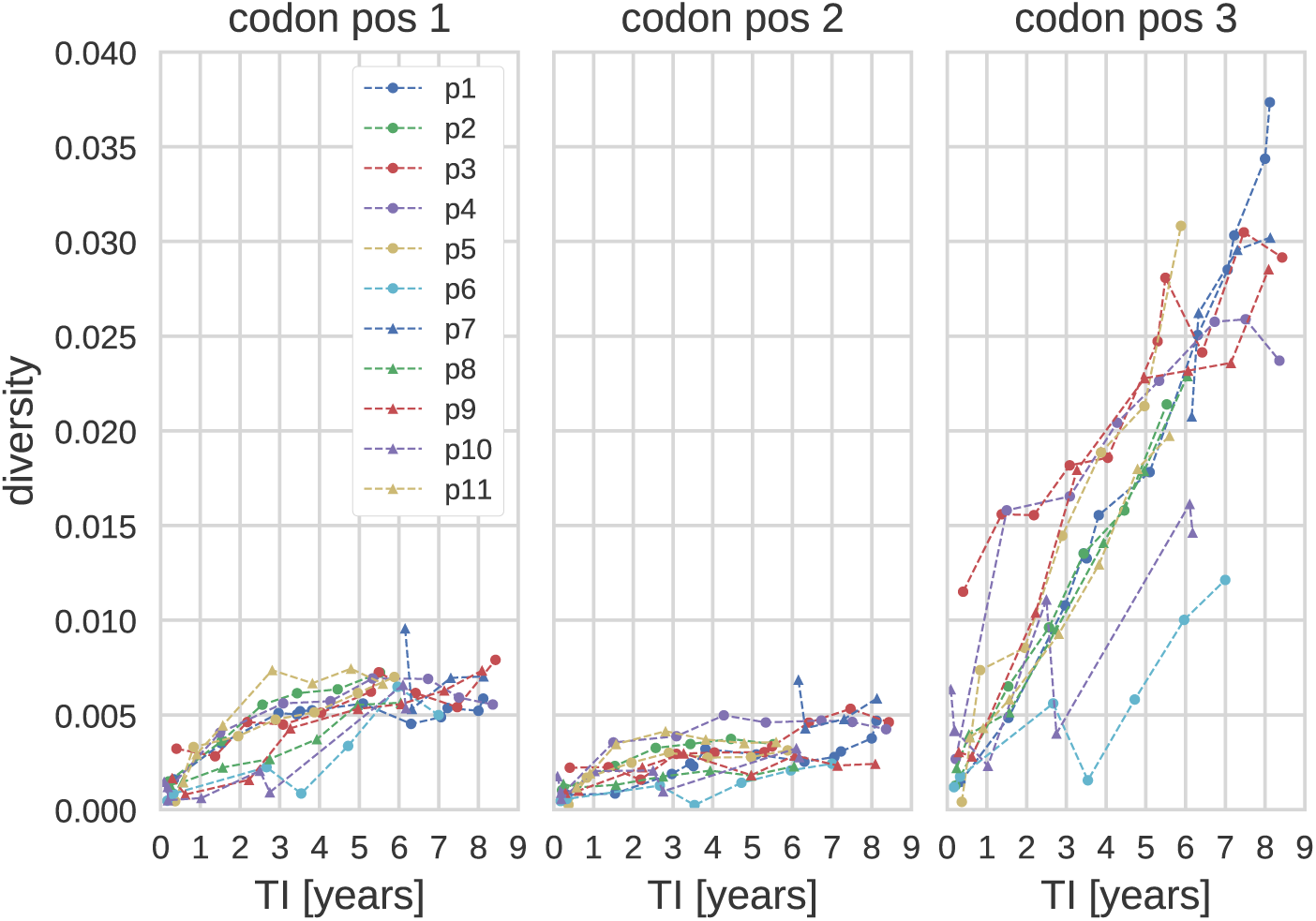
Diversity in pol as a function of the time since infection (TI) and 1st, 2nd and 3rd codon positions. (Genetic region: *pol*, diversity measure: average pairwise distance, *x*_*c*_ = 0.003.)

We quantified the fraction of variation in diversity measures that could be explained by a linear regression of sample date vs. diversity using the coefficient of determination (*r*^2^), see the S3 Fig and the S4 Fig. In all patients for whom early samples were available, a linear regression explained between 70% and 90% of variation if rare iSNVs below 20% population frequency were included, that is the cutoff *x*_*c*_ was below 0.2. The coefficient of determination was much smaller when only iSNVs between 20% and 80% were included, that is the *x*_*c*_ cutoff is larger than 0.2. This decrease is due to increased noise as fewer and fewer sites contribute to the diversity measures.

Furthermore, as seen from S4 Fig, the diversity at 3rd codon positions (at which mutations are mostly synonymous) exhibited higher *r*^2^, whereas the trajectories at the 1st and 2nd codon positions saturated quickly after the infection, Fig. 1. Thus, in the following we limit the analysis mainly to sites in 3rd codon positions (whenever we are dealing with a whole gene, i.e. when the reading frame is known.) However, the results reported below show that inclusion of 1st and 2nd codon positions only has a small deleterious effect on the accuracy of the TI estimates.

### Diversity in *pol* yielded the most accurate TI estimates

Accurate estimation of TI requires averaging diversity across many sites. To investigate which regions of the genome yields the most accurate estimates and how many sites should be averaged, we calculated the average prediction error for averaging windows of different length and in different regions of the genome. In Fig. 2 the mean absolute error (MAE) for the estimated TIs is shown as a function of the genome window position for different window sizes. We found that the most precise TI estimates were obtained for windows with a length of 2000-3000bp. The precision was highest if the window covered the *pol* gene and significantly lower if the window included *env*. Smaller windows contain fewer sites and therefore gave less precise estimates. Larger windows also gave less precise TI estimates because they necessarily include regions in which diversity saturates (such as *env*, S2 Fig) as well as regions that often were sequenced less deeply in our dataset (again *env*, see [24]).

**Fig 2.**
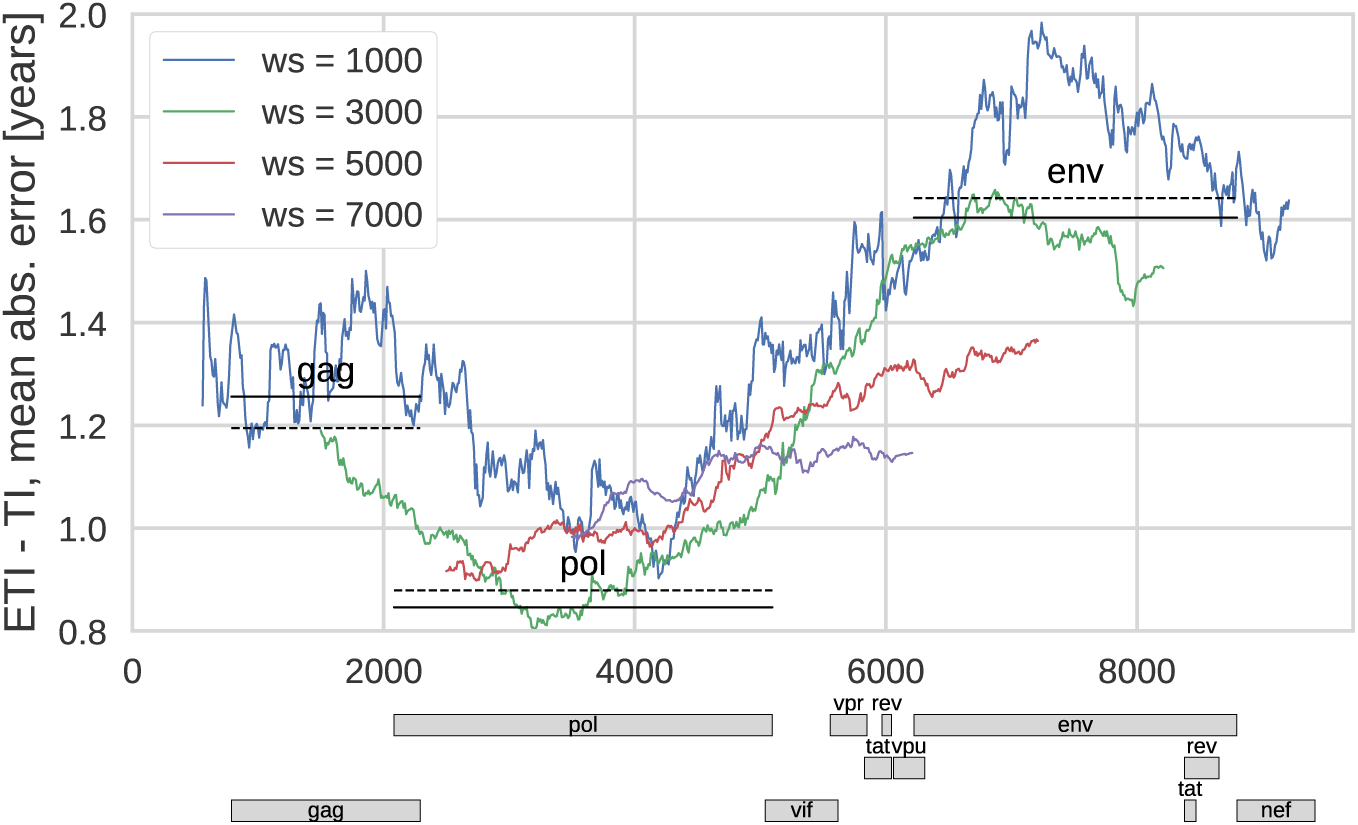
Mean absolute prediction error of TI as a function of position in genome and different sizes of the genome window. (*ws*). Straight solid lines correspond to the error when estimation is based on diversity in the genes *gag*, *pol* or *env*. The dashed lines are analogous estimates using diversity only at 3rd codon position. Diversity measure: average pairwise distance, *x*_*c*_ = 0.003.

The precision of the TI estimates obtained for windows corresponding to particular genes are shown in Fig. 2 by lines indicating the position of the gene and the corresponding average absolute error of the TI estimate. The dashed lines correspond to estimates using only 3rd codon positions of the corresponding gene.

### Accuracy increased with higher sequencing resolution

Genetic diversity in NGS data can be quantified by different measures and we investigated the performance of three related measures – average pairwise distance, site entropy, and fraction of polymorphic sites. As discussed above, these measures put different emphasis on iSNVs at different frequency. Even though the average pairwise distance and site entropy formally do not require a cutoff on minority iSNVs, in practice a cutoff is necessary to remove low-level experimental errors. We therefore introduced an iSNV cutoff in calculating the diversity based on average pairwise distance and site entropy, *x*_*c*_, taking into account only the data with iSNVs above *x*_*c*_. For diversity measures based on polymorphic sites, the cutoff value corresponds to the iSNV level from which the site is considered polymorphic. This emulates ambiguous base calls by Sanger sequencers, which (at best) can identify minority iSNVs above a threshold of around 25% [25–27].

Given the high ability of NGS to detect low level iSNVs one can consider the dependence of the TI estimation on the cutoff value, as shown for all three diversity measures in Fig. 3. All three diversity measures performed equally well and increasingly better at cutoff values *x*_*c*_ down to around 10%. This indicates that diversity in NGS data allows more accurate estimates of TI than ambiguous base calls in Sanger sequences and that this primarily is due to better sensitivity and accuracy in detection of low level iSNVs. For cutoffs below 10% the error of TI estimates based on counting polymorphic sites was greater than those based on the other two measures since rare iSNVs which are sensitive to sequencing errors and amplification biases contribute as much as common iSNVs. Indeed, at *x*_*c*_ → 0 this measure includes all sites and becomes completely insensitive to differences in iSNV levels. The other two measures do not suffer from this problem, because they put different weights on sites with different iSNV levels. Therefore the prediction errors for estimates based 3rd codon positions and average pairwise distance or site entropy continued to decrease all the way down to *x*_*c*_ = 0.003, which represents the cutoff for sequencing errors for our NGS method [24].

**Fig 3.**
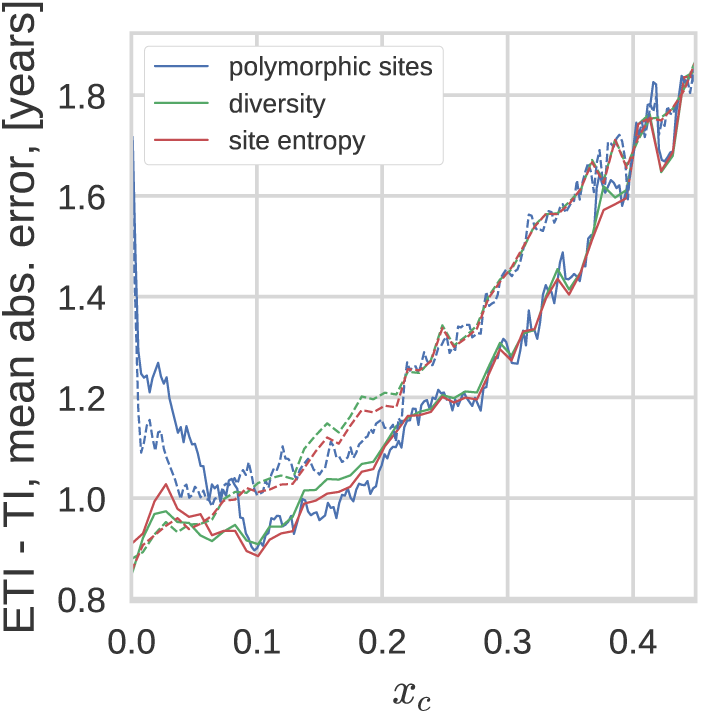
Mean absolute error as a function of the low-frequency cutoff. (*x*_*c*_). Different diversity measures perform very similarly when the cutoff *x*_*c*_ is greater than approximately 10%. Average pairwise distance and entropy outperform fraction of polymorphic sites for low *x*_*c*_. This graph is based on diversity in *pol*. Solid lines correspond to using all sites, dashed lines to diversity at 3rd codon positions.

The noticeably non-monotonic behavior of the predictions based on average pairwise distance and site entropy when all codon positions are taken into account is due to the saturation of 1st and 2nd codon positions (i.e. non-synonymous) diversity. The 1st and 2nd codon positions tend to be conserved since they result in mostly non-synonymous mutations. Depending on the fitness cost associated with the mutation, diversity at these sites saturates at different levels [37]. As the threshold *x*_*c*_ is lowered, sites with higher and higher fitness costs contribute to the diversity measures and the effect of the saturation behavior becomes more pronounced. Note that the non-monotonic dependence is not consistently reproduced in other genes (see supplementary S5 Fig). Similarly, the fact that diversity measures based on all codon positions perform somewhat better at *x*_*c*_ *>* 10% was not consistently reproduced in other genes. In *gag*, diversity at 3rd positions was better at estimating TI for most *x*_*c*_ values (S5 Fig).

In the following analyses we opted for using the average pairwise distance measure, taking into account only the diversity at the 3rd codon positions, with low (*x*_*c*_ = 0.003) iSNV cutoff. Average pairwise distance was chosen because the results were virtually indistinguishable from those produced using site entropy, but easier to calculate and interpret. Since 1st and 2nd codon positions contribute very little time-dependent diversity and are affected by purifying selection and selective sweeps, we recommend to restrict the diversity measures to 3rd codon positions.

### Distribution of prediction errors

The results above indicated that more than 50% of the estimated TIs fell within a window of about one year centered at the actual TI. Fig. 4 (Left) shows a more direct analysis of the distribution of TI prediction errors. The distribution is tightly peaked around zero, but has a left tail corresponding to samples estimated to have been obtained shorter after infection than they actually were drawn, i.e. diversity being lower than expected. Most of these samples were from p6, who throughout infection had lower diversity than other patients. In addition to biological reasons for low diversity, amplification problems and low RNA template numbers (i.e. low virus levels) can explain underestimation of diversity.

**Fig 4.**
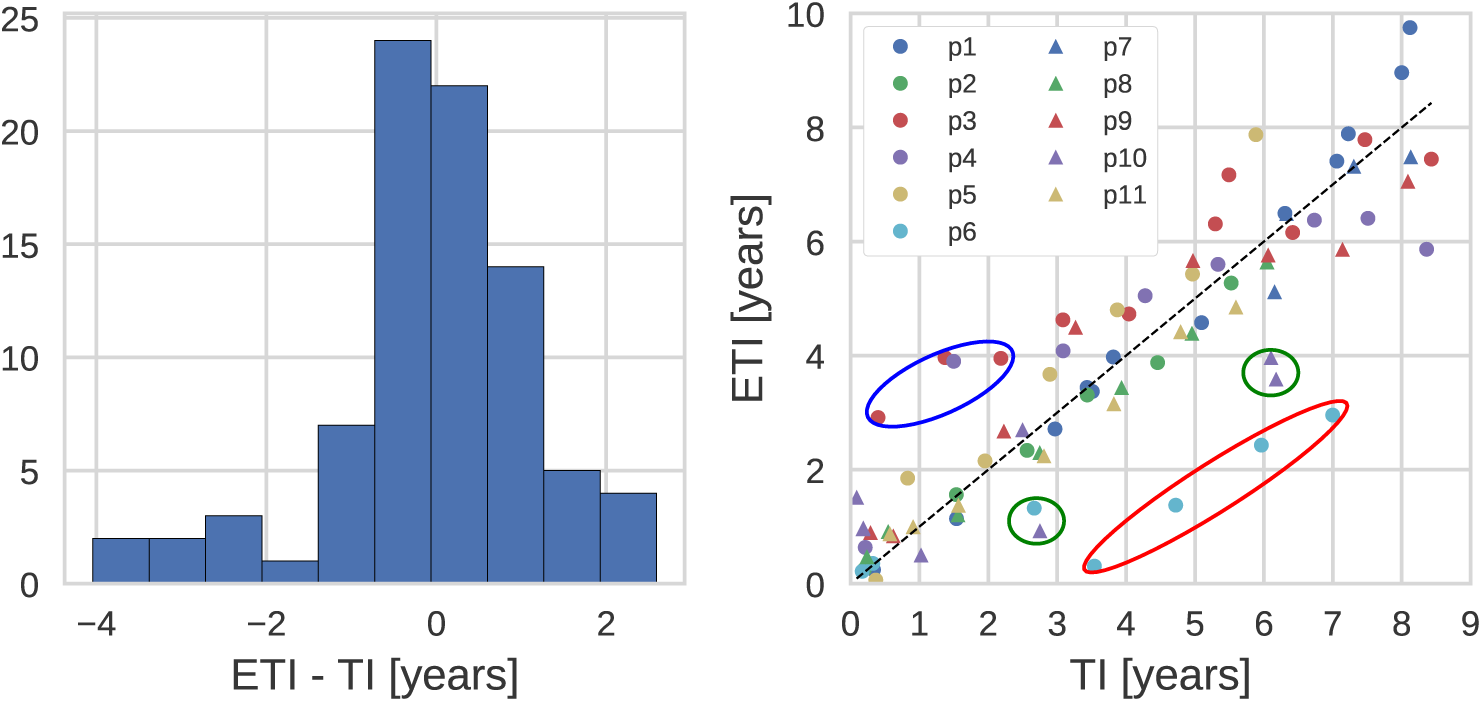
(Left) Distribution of the estimation error. (Right) Estimated time of infection (ETI) versus actual time of infection (TI). (Genetic region: 3rd codon positions in *pol*, diversity measure: average pairwise distance. The encircled outliers are discussed in the text.)

Some samples were estimated to have been drawn later after infection than the true duration of infection. In particular early samples from p10 and p3 had markedly higher diversity than expected. For both patients, we have evidence that their infections were established by more than one virion resulting in carry-over of diversity from the donor. This excess diversity gradually decreased in p10 and, somewhat slower, in p3.

Next, we investigated how the prediction error depended on the time since infection. Fig. 5 shows the average absolute error of the estimated TI versus the true TI, averaged over *n* = 25 consecutive data points. This average error (see for details S2 Appendix) was surprisingly stable over TIs, and only increased slightly from around 0.6 years to around 1.1 years as the age of infection increased from 6 months to 6 years. This increase can be attributed to bigger statistical fluctuations of diversity later after infection due to factors such as variations in the rate of diversification or differences in number and strength of selective sweeps that reduce diversity.

**Fig 5.**
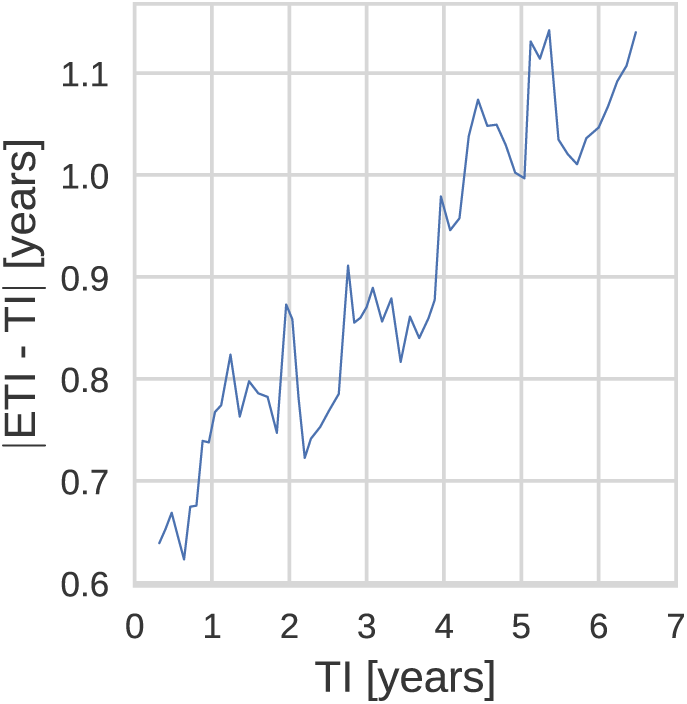
Estimation error dependence on time of infection (TI): TI and |ETI-TI| averaged over *n* = 25 adjacent points. (Genetic region: 3rd codon positions in *pol,* diversity measure: average pairwise distance.)

### Recommended regression coefficients

In Table 2 we list the values of slope and intercept that can be used to estimate the infection date from the known diversity calculated as average pairwise distances for 3rd codon positions in *pol* gene. The supplementary materials contain similar tables for the two other diversity measures (S1 Table and S2 Table), as well as for the case when all codon positions are taken into account (S3 Table, S4 Table and S5 Table). As the iSNV resolution may vary between different sequencing methods and facilities, we list the values of the parameters for different cutoffs, implying that all the frequencies below the cutoff value are set to zero and the corresponding sites do not contribute to the diversity measure. Note that the slope (and intercept) increases with increasing iSNV cutoffs because fewer and fewer sites contribute to diversity. In addition to the two parameter model, we also investigated the performance of a model with the slope as the single parameter, i.e. no intercept (*t*_0_ = 0). This model has a slightly higher absolute prediction error. However, for low values of the cutoff *x*_*c*_ *≈* 5% these models agree and for cutoffs below 20% the two models perform equally well.

**Table 2.**
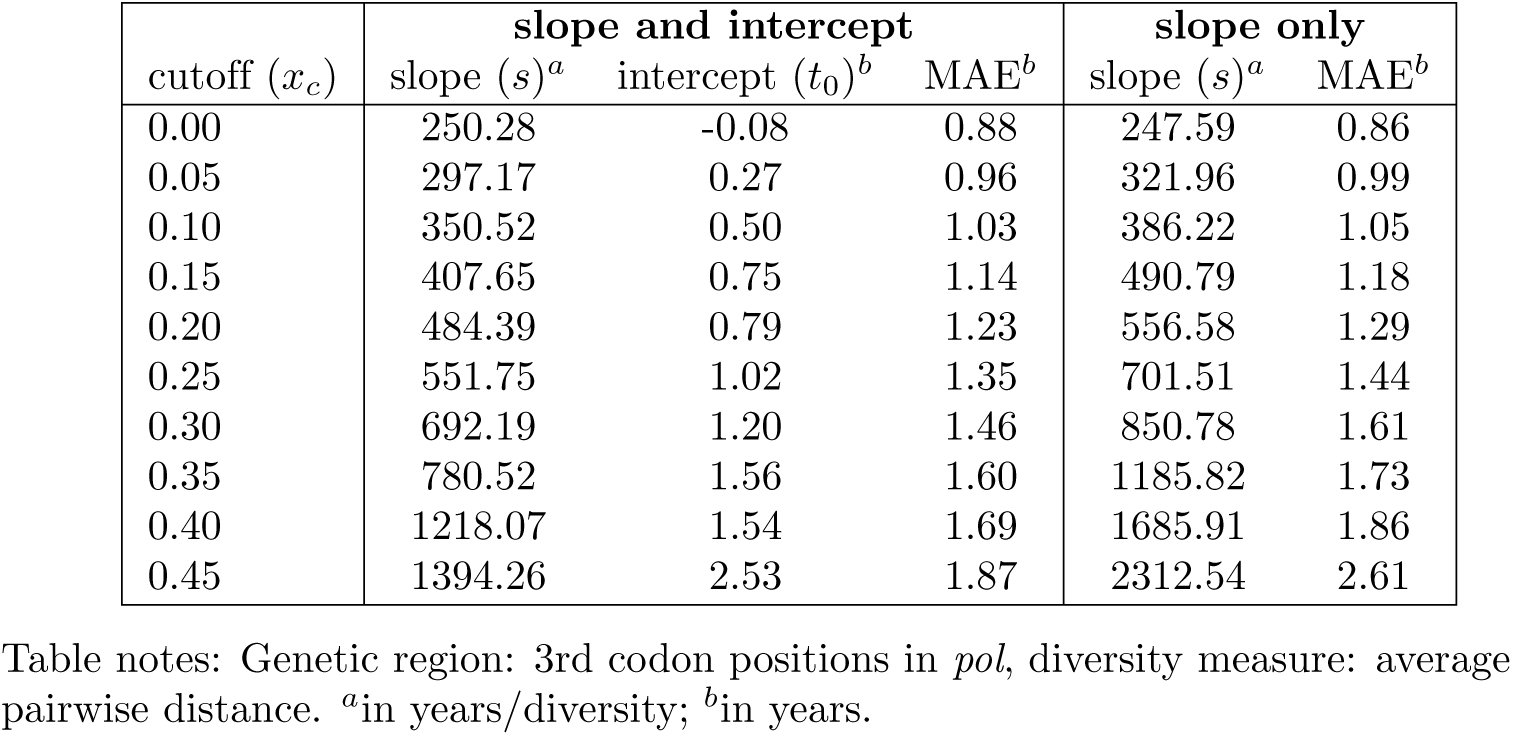
Recommended slope and intercept values depending on the cutoff.

| cutoff ($x_c$) | slope and intercept | | | slope only | |
| --- | --- | --- | --- | --- | --- |
| | slope ($s$)<sup>a</sup> | intercept ($t_0$)<sup>b</sup> | MAE<sup>b</sup> | slope ($s$)<sup>a</sup> | MAE<sup>b</sup> |
| 0.00 | 250.28 | -0.08 | 0.88 | 247.59 | 0.86 |
| 0.05 | 297.17 | 0.27 | 0.96 | 321.96 | 0.99 |
| 0.10 | 350.52 | 0.50 | 1.03 | 386.22 | 1.05 |
| 0.15 | 407.65 | 0.75 | 1.14 | 490.79 | 1.18 |
| 0.20 | 484.39 | 0.79 | 1.23 | 556.58 | 1.29 |
| 0.25 | 551.75 | 1.02 | 1.35 | 701.51 | 1.44 |
| 0.30 | 692.19 | 1.20 | 1.46 | 850.78 | 1.61 |
| 0.35 | 780.52 | 1.56 | 1.60 | 1185.82 | 1.73 |
| 0.40 | 1218.07 | 1.54 | 1.69 | 1685.91 | 1.86 |
| 0.45 | 1394.26 | 2.53 | 1.87 | 2312.54 | 2.61 |

Table notes: Genetic region: 3rd codon positions in *pol*, diversity measure: average pairwise distance. <sup>a</sup>in years/diversity; <sup>b</sup>in years.

In order to make our data and the method of TI inference more accessible for practical use, we added a web application to the web site containing the processed patient data, accessible at hiv.biozentrum.unibas.ch/ETI/. Given a diversity value (determined according to Eq. (2)), the web application allows to determine the time since infection for an user-selected iSNV cutoff value (*x*_*c*_) and genetic region, along with the appropriate error estimates. The results are presented in accessible graphical form, but also as a slope and intercept pair, equivalent to Table. 2. The user can specify whether the diversity was calculated using all codon positions or only the 1st, 2nd, or 3rd. Specifying a codon position is only supported when the region used to evaluate diversity is fully contained in one gene.

### Validation of the regression coefficients

We validated the regression coefficients on a dataset from 31 patients with known infection dates and NGS data available at two time points. The distribution of the diversity values for these patients closely resembles that for our training dataset of 11 patients (see S8 Fig and S9 Fig).

We inferred the time since infection for the 31 validation patients using the regression coefficients obtained for the eleven patients of the training set; the results are summarized in Fig. 6, which also shows (in gray) the data points of the training set (same as in Fig. 4). The new data exhibit the same behavior as the training set: the TI estimates are centered around the true TI and the accuracy of the estimate is about one year, as can be seen from the histogram in the left panel of Fig. 6. In order to make the histograms for the training and validation data comparable, we included for the former only the points of the infection time less than 5 years.

**Fig 6.**
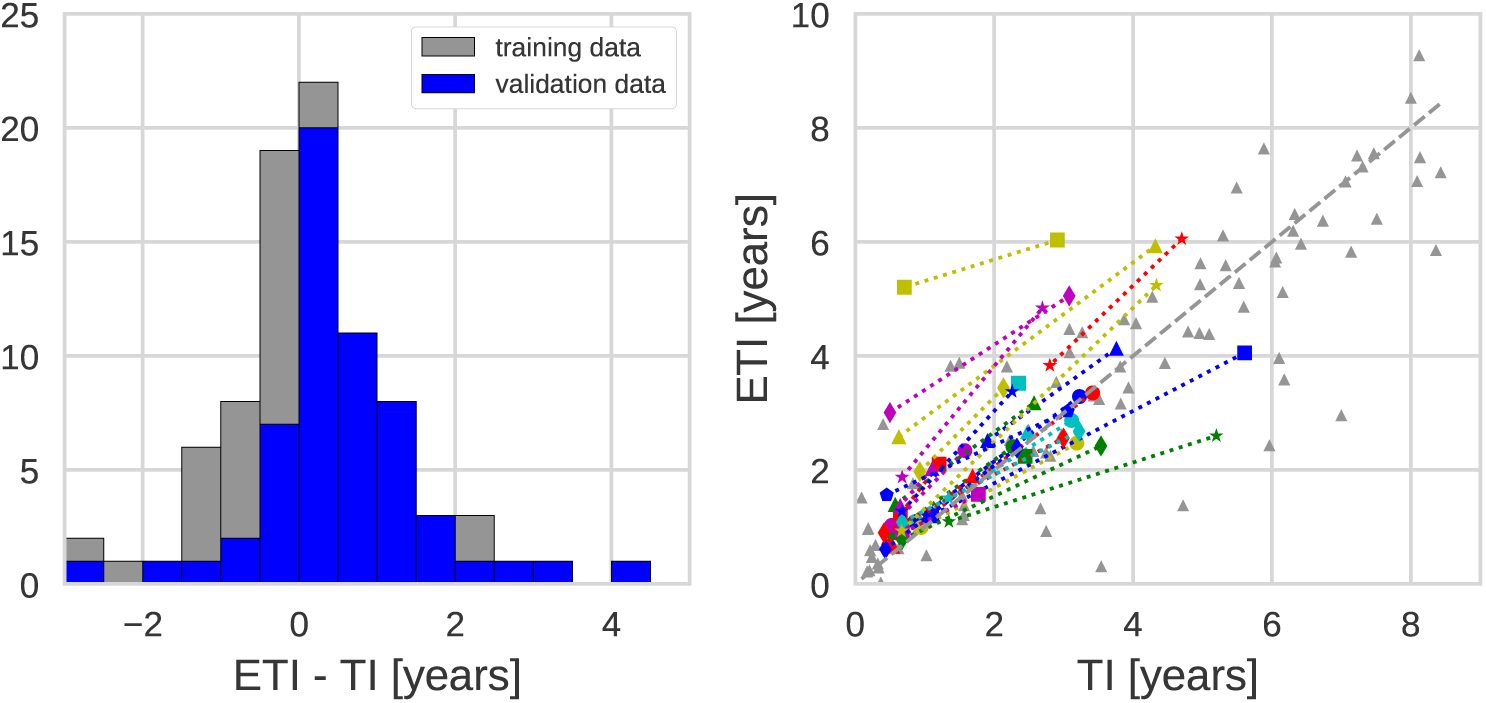
(Left) Distribution of the estimation error. (Right) Estimated time of infection (ETI) versus actual time of infection (TI). Displayed for the training and the validation data sets. (Genetic region: 3rd codon positions in *pol*, diversity measure: average pairwise distance, *x*_*c*_ = 0.003. Connected data points belong to the same patient.)

Some outliers are present also in validation data, particularly data points with overestimated TIs, i.e. having higher diversity than expected. As mentioned above, these data points probably often represent infections established by more than one founder virus. One patient had substantial overestimation of time since infection, which might be due to superinfection from different donors rather than multiple founders from a single donor. However, it should be stressed that almost all outliers are still within +/− 2 years from the true TI. Neither overestimation nor underestimation of TI was clearly related to genetic subtype of the virus, transmission route, virus levels or template numbers (S11 Fig and S12 Fig).

## Discussion

Many newly diagnosed HIV-1 patients have an infection of unknown duration. This is problematic because accurate estimation of the time since infection (TI) is essential for understanding important aspects of HIV-1 epidemiology such as incidence, proportion of undiagnosed patients and late presentation. Most previous methods are suboptimal because they only categorize patients as being recently or long-term infected and/or are imprecise. Here, we show that genetic diversity calculated from NGS data enables fairly accurate estimation of TI, even many years after infection. We also show that NGS is superior to Sanger sequencing because inclusion of minority iSNVs below the Sanger detection limit (around 25%) substantially improves the accuracy of the TI estimates.

We investigated how the TI estimates were influenced by sequence length, genome region, codon position, iSNV cutoff and type of distance measure. We found that the most precise estimates of TI were obtained using average pairwise distance or site entropy based on 3rd codon positions in the *pol* gene. For these data viral diversity increases approximately linearly during at least 8 years after infection, which allows estimation of TI during this time period. The accuracy of the TI estimate was approximately +/− 1 year in long-term infections, and slightly better during the first year of infection. The *env* gene was less suitable than *pol* for estimating TI, especially if longer time had elapsed since infection. This is because frequent selective sweeps in *env* continuously remove diversity and this effect becomes increasingly evident with increasing time since infection [17, 24]. This explains why the most accurate results were obtained using 2000-3000 base pair long sequences covering *pol*, while omitting *env*. We found that most of temporal signal came from 3rd codon positions (at which most mutations are synonymous) and that omission of 1st and 2nd codon positions slightly improved TI estimates for iSNV cutoffs (*x*_*c*_) below 10%, i.e. when the full potential of NGS was utilized. Diversity measures based on average pairwise distance and site entropy outperformed the measure based on fraction of polymorphic sites at low iSNV cutoffs.

Based on our results we make the following recommendations for TI estimation based on HIV NGS data; average pairwise distance on 3rd codon positions in *pol* sequences with the lowest possible cutoff for iSNVs (in our case 0.3%). Importantly, we have validated our recommendations by applying them to a validation dataset consisting of NGS data from 31 additional patients with known infection times. The accuracy of TI estimation for the validation data was as accurate and precise as for the training data, which confirms the applicability of our method for broader clinical and epidemiological use. For convenience we provide a table that translates viral diversity into TI as well as a web application that estimates TI for user-defined regions of the HIV-1 genome.

Even though we primarily have focused on estimation of TI in individual patients, our method can be applied to estimation of incidence in populations. Many methods for HIV-1 incidence estimation in populations have been based on biomarkers that classify patients as recently or long-term infected (e.g. the BED and LAg assays). [1–7]. Such binary classification can be done based on NGS data; if the diversity is less than a specified cutoff value *D*_*cr*_ infection is classified as recent, and otherwise as long-term. The cutoff between recent and long-term infections can be chosen by the investigator using Table 2 or the website. For instance a diversity of 0.0021 for 3rd bases on *pol* corresponds to 180 days since infection. However, we have not fully determined two test properties that are required for most binary incidence estimators; the mean duration of recent infection (MDRI) and the false-recent rate (FRR) [38]. The NGS data can also be used to directly model HIV-1 incidence based on TIs [39, 40].

Our study has some limitations. Ideally we should have studied a larger and more diverse set of training and validation patients. However, patients with known time of infection, long followup without therapy and suitable biobank samples are rare. Today it would be unethical to delay start of ART. Thus, our training dataset consisted of patients who were diagnosed between 1990 and 2003 and retrospectively identified and investigated using stored biobank samples [24]. The patients in the validation dataset were diagnosed between 2003 and 2010, and as a consequence had shorter followup without ART. In the training dataset 9 of 11 patients were MSM infected with subtype-B virus. The validation patients had a more diverse distribution with 11 patients infected by other routes than MSM and 16 patients infected with non-B-subtypes. The precision in the TI estimate was similar for patients with MSM and non-MSM as well as subtype B and non-B infections, which again suggests that our regression coefficients can be broadly applied (S11 Fig and S12 Fig).

Another limitation is that the true (i.e. “known”) TI was estimated from laboratory and clinical data and therefore has an error that we have not considered because it is difficult to estimate. However, such a measurement error, which surely exists, will reduce the accuracy at which TI can be estimated and assuming zero measurement error for the “true” TI is therefore conservative.

A potential problem with NGS data is incomplete sampling of virus diversity in samples with low virus levels. If the sequencing library is dominated by a few template molecules the TI estimate might be erroneously short. A related problem is due to the reduced ability of NGS to correctly estimate TI in infections that were founded by multiple virions. Two of our 11 training patients showed evidence of such multiple infections. Clear overestimation of TI was also observed in one of the 31 validation patients. It has been reported that HIV-1 infection is founded by more than one virion in around 40% of MSM and around 20% of heterosexuals, whereas superinfection (from different donors) is more rare [41, 42]. In view of the fact that most of our study patients were MSM, it is surprising that serious overestimation of TI was not observed more often. There are two possible explanations for this. Firstly, TI will only be overestimated if the multiple founders are sufficiently diverse. Secondly, the overestimation of TI appeared to diminished over time in the two training patients, which may happen if excess diversity is removed because favored iSNVs are selected for over time [24]. Even though our method has limitations with multiple founders and superinfection, a diversity value that exceeds the upper 95% confidence value of the diversity that expected 10 years or more after infection should be treated with great caution, because it may be due to multiple founders and/or superinfection. Furthermore, we plan to investigate estimation of TI can be improved by combining virus diversity determined NGS with other biomarkers, such as BED, LAg avidity, CD4 and virus levels, in a multiple assay algorithm.

Finally, NGS is not yet part of routine diagnostics for HIV resistance. However, in the coming years NGS can be expected to replace Sanger sequencing for clinical HIV-1 resistance testing, which is recommended for all newly diagnosed patients (in developed countries). Thus, while our method for estimating TI from NGS data currently requires extra laboratory work, NGS data is likely to become increasingly available as part of routine HIV-1 care, which will increase the utility of our method.

### Conclusion

In conclusion, we show that sequence diversity determined by NGS can be used to estimate time since HIV-1 infection with a precision that is better than most alternative biomarkers. Importantly, TI can be estimated many years after infection, whereas most alternative methods only categorize patients as being recently or long-term infected or are less precise. We found that TI was most accurately estimated using 3rd codon positions in *pol* sequences with a *x*_*c*_ = 0.003 cutoff for iSNVs and that average pairwise distances was the preferred distance measure. Samples with low virus levels and infections established by multiple virions can give rise to misleading levels of virus diversity. Algorithms based on NGS diversity in combinations with other biomarkers may prove very useful.

## Supporting information

**S1 Fig. Diversity in *gag* as a function of the time since infection (TI**). (Diversity measure: average pairwise distance, *x*_*c*_ = 0.003.)

**S2 Fig. Diversity in *env* as a function of the time since infection (TI).** (Diversity measure: average pairwise distance, *x*_*c*_ = 0.003.)

**S3 Fig. Coefficient of determination for different diversity measures, including all sites**. (Genetic region: *pol*.)

**S4 Fig. Coefficient of determination for average pairwise distance (diversity), by codon position**. (Genetic region: *pol*.)

**S5 Fig. Mean absolute error as a function of the low-frequency cutoff** (*x*_*c*_). Different diversity measures perform very similarly when the cutoff *x*_*c*_ is large. Average pairwise distance and entropy outperform fraction of polymorphic sites for low *x*_*c*_. This graph is based on diversity in *gag* (left) and *env* (Right). Solid lines correspond to using all sites, dashed - only 3rd codon positions.

**S6 Fig. Distribution of slopes (*s*) and intercepts (*t*_0_) (bootstrapped over the patients of the training dataset**). (Genetic region: 3rd codon positions in *pol*, diversity measure: average pairwise distance, *x*_*c*_ = 0.003.)

**S7 Fig. Dependence of the slope and intercept in the cutoff.** (Genetic region: 3rd codon positions in *pol*, diversity measure: average pairwise distance.)

**S8 Fig. Diversity as a function of the time since infection (TI) and 1st, 2nd and 3rd codon positions. Displayed for the training and the validation datasets**. (Genetic region: *gag*, diversity measure: average pairwise distance, *x*_*c*_ = 0.003.)

**S9 Fig. Diversity as a function of the time since infection (TI) and 1st, 2nd and 3rd codon positions. Displayed for the training and the validation datasets**. (Genetic region: *gag*, diversity measure: average pairwise distance, *x*_*c*_ = 0.003.)

**S10 Fig. (Left) Distribution of the estimation error. (Right) Estimated time of infection (ETI) versus actual time of infection (TI). Displayed for the training and the validation data sets**. (Genetic region: 3rd codon positions in *gag*, diversity measure: average pairwise distance, *x*_*c*_ = 0.003. The encircled outliers are discussed in the text.)

**S11 Fig. Estimated time of infection (ETI) versus actual time of infection (TI). Displayed for the training and the validation data sets. Legend shows the subtype (top) and transmission route (bottom) for the validation dataset patients**. (Genetic region: 3rd codon positions in *pol*, diversity measure: average pairwise distance, *x*_*c*_ = 0.003.)

**S12 Fig. Estimated time of infection (ETI) versus actual time of infection (TI). Displayed for the training and the validation data sets. Legend shows the number of templates (top) and the dilutions (bottom) for the validation dataset.** (Genetic region: 3rd codon positions in *pol*, diversity measure: average pairwise distance, *x*_*c*_ = 0.003.)

### S1 Appendix Linear fitting procedures

#### S2 Appendix Moving average

**S1 Table Recommended slope and intercept values depending on the cutoff** (Genetic region: 3rd codon positions in *pol*, diversity measure: average number of polymorphic sites. ^*a*^in years/diversity; ^*b*^in years.)

**S2 Table Recommended slope and intercept values depending on the cutoff.** (Genetic region: 3rd codon positions in *pol*, diversity measure: average site entropy ^*a*^in years/diversity; ^*b*^in years.)

**S3 Table Recommended slope and intercept values depending on the cutoff.** (Genetic region: all sites in *pol*, diversity measure: average pairwise distance. ^*a*^in years/diversity; ^*b*^in years.)

**S4 Table Recommended slope and intercept values depending on the cutoff.** (Genetic region: all sites in *pol*, diversity measure: average number of polymorphic sites. ^*a*^in years/diversity; ^*b*^in years.)

**S5 Table Recommended slope and intercept values depending on the cutoff.** (Genetic region: all sites in *pol*, diversity measure: average site entropy. ^*a*^inyears/diversity; ^*b*^in years.)

**S6 Table Expanded summary of patient characteristics (training dataset).** See also S1 Text

**S7 Table Expanded summary of patient characteristics (validation dataset).**

**S1 Text Supporting text for S6 Table.**

## Acknowledgments

We would like to express our gratitude to the study participants and to Johanna Brodin, Fabio Zanini, Lina Thebo, Christa Lanz and Göran Bratt for important contributions in the generation of the published data that was analyzed in this study.

## References

1. Le Vu S, Pillonel J, Semaille C, Bernillon P, Le Strat Y, Meyer L, et al. Principles and uses of HIV incidence estimation from recent infection testing–a review. Euro Surveill. 2008;13(36):537–545.

2. Incidence Assay Critical Path Working Group. More and better information to tackle HIV epidemics: towards improved HIV incidence assays. PLoS Med. 2011;8(6):e1001045.

3. World Health Organization. When and how to use assays for recent infection to estimate HIV incidence at a population level. 2011;.

4. Busch MP, Pilcher CD, Mastro TD, Kaldor J, Vercauteren G, Rodriguez W, et al. Beyond detuning: 10 years of progress and new challenges in the development and application of assays for HIV incidence estimation. Aids. 2010;24(18):2763–2771.

5. Murphy G, Parry J. Assays for the detection of recent infections with human immunodeficiency virus type 1. Euro Surveill. 2008;13(36):314–320.

6. Park SY, Love TM, Nelson J, Thurston SW, Perelson AS, Lee HY. Designing a genome-based HIV incidence assay with high sensitivity and specificity. AIDS (London, England). 2011;25(16):F13–F19.

7. Moyo S, Vandormael A, Wilkinson E, Engelbrecht S, Gaseitsiwe S, Kotokwe KP, et al. Analysis of Viral Diversity in Relation to the Recency of HIV-1C Infection in Botswana. PLoS One. 2016;11(8):e0160649.

8. Guy R, Gold J, Calleja JMG, Kim AA, Parekh B, Busch M, et al. Accuracy of serological assays for detection of recent infection with HIV and estimation of population incidence: a systematic review. The Lancet infectious diseases. 2009;9(12):747–759.

9. Bärnighausen T, McWalter TA, Rosner Z, Newell ML, Welte A. Review Article: HIV Incidence Estimation Using the BED Capture Enzyme Immunoassay: Systematic Review and Sensitivity Analysis. Epidemiology. 2010; p. 685–697.

10. Wei X, Liu X, Dobbs T, Kuehl D, Nkengasong JN, Hu DJ, et al. development of two avidity-based assays to detect recent HIV type 1 seroconversion using a multisubtype gp41 recombinant protein. Aids research and human retroviruses. 2010;26(1):61–71.

11. Lazarus J, Jürgens R, Weait M, Phillips A, Hows J, Gatell J, et al. Overcoming obstacles to late presentation for HIV infection in Europe. HIV medicine. 2011;12(4):246–249.

12. Sabin CA, Schwenk A, Johnson MA, Gazzard B, Fisher M, Walsh J, et al. Late diagnosis in the HAART era: proposed common definitions and associations with mortality. Aids (London, England). 2010;24(5):723–727.

13. Goujard C, Bonarek M, Meyer L, Bonnet F, Chaix ML, Deveau C, et al. CD4 cell count and HIV DNA level are independent predictors of disease progression after primary HIV type 1 infection in untreated patients. Clinical Infectious Diseases. 2006;42(5):709–715.

14. Lodi S, Phillips A, Touloumi G, Geskus R, Meyer L, Thiébaut R, et al. Time from human immunodeficiency virus seroconversion to reaching CD4+ cell count thresholds 200, 350, and 500 cells/mm3: assessment of need following changes in treatment guidelines. Clinical infectious diseases. 2011;53(8):817–825.

15. Lodi S, Guiguet M, Costagliola D, Fisher M, de Luca A, Porter K. Kaposi sarcoma incidence and survival among HIV-infected homosexual men after HIV seroconversion. Journal of the National Cancer Institute. 2010;102(11):784–792.

16. Minga A, Lewden C, Gabillard D, Bomisso G, Toni Td, Emième A, et al. CD4 eligibility thresholds: an analysis of the time to antiretroviral treatment in West African HIV-1 seroconverters. Aids (London, England). 2011;25(6):819–823.

17. Shankarappa R, Margolick JB, Gange SJ, Rodrigo AG, Upchurch D, Farzadegan H, et al. Consistent viral evolutionary changes associated with the progression of human immunodeficiency virus type 1 infection. Journal of virology. 1999;73(12):10489–10502.

18. Kouyos R, von Wyl V, Yerly S, Böni J, Rieder P, Joos B, et al. Ambiguous nucleotide calls from population-based sequencing of HIV-1 are a marker for viral diversity and the age of infection. Clinical infectious diseases: an official publication of the Infectious Diseases Society of America. 2011;52(4):532–539.

19. Ragonnet-Cronin M, Aris-Brosou S, Joanisse I, Merks H, Vallée D, Caminiti K, et al. Genetic Diversity as a Marker for Timing Infection in HIV-Infected Patients: Evaluation of a 6-Month Window and Comparison With BED. Journal of Infectious Diseases. 2012;206(5):756–764.

20. Andersson E, Shao W, Bontell I, Cham F, Wondwossen A, Morris L, et al. Evaluation of sequence ambiguities of the HIV-1 pol gene as a method to identify recent HIV-1 infection in transmitted drug resistance surveys. Infection, Genetics and Evolution. 2013;18:125–131.

21. Meixenberger K, Hauser A, Jansen K, Yousef KP, Fiedler S, von Kleist M, et al. Assessment of ambiguous base calls in HIV-1 pol population sequences as a biomarker for identification of recent infections in HIV-1 incidence studies. Journal of clinical microbiology. 2014;52(8):2977–2983.

22. Allam O, Samarani S, Ahmad A. Hammering out HIV-1 incidence with Hamming distance. Aids. 2011;25(16):2047–2048.

23. Cousins MM, Konikoff J, Laeyendecker O, Celum C, Buchbinder SP, Seage GR, et al. HIV diversity as a biomarker for HIV incidence estimation: including a high-resolution melting diversity assay in a multiassay algorithm. Journal of clinical microbiology. 2014;52(1):115–121.

24. Zanini F, Brodin J, Thebo L, Lanz C, Bratt G, Albert J, et al. Population genomics of intrapatient HIV-1 evolution. Elife Sciences. 2016;4:e11282. doi:10.7554/eLife.11282.

25. Schuurman R, Brambilla D, de Groot T, Huang D, Land S, Bremer J, et al. Underestimation of HIV type 1 drug resistance mutations: results from the ENVA-2 genotyping proficiency program. Aids research and human retroviruses. 2002;18(4):243–248.

26. Parkin N, Bremer J, Bertagnolio S. Genotyping external quality assurance in the World Health Organization HIV drug resistance laboratory network during 2007–2010. Clinical infectious diseases. 2012;54(Suppl 4):S266–S272.

27. Land S, Cunningham P, Zhou J, Frost K, Katzenstein D, Kantor R, et al. TREAT Asia Quality Assessment Scheme (TAQAS) to standardize the outcome of HIV genotypic resistance testing in a group of Asian laboratories. Journal of virological methods. 2009;159(2):185–193.

28. Fiebig EW, Wright DJ, Rawal BD, Garrett PE, Schumacher RT, Peddada L, et al. Dynamics of HIV viremia and antibody seroconversion in plasma donors: implications for diagnosis and staging of primary HIV infection. Aids. 2003;17(13):1871–1879.

29. Gaines H, von Sydow M, Pehrson PO, Lundbegh P. Clinical picture of primary HIV infection presenting as a glandular-fever-like illness. BMJ. 1988;297(6660):1363–1368.

30. Cohen MS, Shaw GM, McMichael AJ, Haynes BF. Acute HIV-1 infection. New England Journal of Medicine. 2011;364(20):1943–1954.

31. Skar H, Albert J, Leitner T. Towards estimation of HIV-1 date of infection: a time-continuous IgG-model shows that seroconversion does not occur at the midpoint between negative and positive tests. PLoS One. 2013;8(4):e60906.

32. Zanini F, Brodin J, Albert J, Neher RA. Error rates, PCR recombination, and sampling depth in HIV-1 Whole Genome Deep Sequencing. Virus research. 2016;doi:10.1016/j.virusres.2016.12.009.

33. Cock PJA, Antao T, Chang JT, Chapman BA, Cox CJ, Dalke A, et al. Biopython: freely available Python tools for computational molecular biology and. Bioinformatics. 2009;25(11):1422–1423. doi:10.1093/bioinformatics/btp163.

34. van der Walt S, Colbert SC, Varoquaux G. The NumPy Array: A Structure for Efficient Numerical Computation. Computing in Science Engineering. 2011;13(2):22–30. doi:10.1109/MCSE.2011.37.

35. Hunter JD. Matplotlib: A 2D graphics environment. Computing In Science & Engineering. 2007;9(3):90–95.

36. Nei M, Li WH. Mathematical model for studying genetic variation in terms of restriction endonucleases. Proceedings of the National Academy of Sciences. 1979;76(10):5269–5273.

37. Zanini F, Puller V, Brodin J, Albert J, Neher RA. In vivo mutation rates and the landscape of fitness costs of HIV-1. Virus. 2017;3:1.

38. Kassanjee R, Pilcher CD, Keating SM, Facente SN, McKinney E, Price MA, et al. Independent assessment of candidate HIV incidence assays on specimens in the CEPHIA repository. Aids (London, England). 2014;28(16):2439.

39. Sommen C, Commenges D, Vu SL, Meyer L, Alioum A. Estimation of the distribution of infection times using longitudinal serological markers of HIV: implications for the estimation of HIV incidence. Biometrics. 2011;67(2):467–475.

40. Romero-Severson EO, Lee Petrie C, Ionides E, Albert J, Leitner T. Trends of HIV-1 incidence with credible intervals in Sweden 2002–09 reconstructed using a dynamic model of within-patient IgG growth. International journal of epidemiology. 2015;44(3):998–1006.

41. Li H, Bar KJ, Wang S, Decker JM, Chen Y, Sun C, et al. High multiplicity infection by HIV-1 in men who have sex with men. PLoS pathogens. 2010;6(5):e1000890.

42. Redd AD, Quinn TC, Tobian AA. Frequency and implications of HIV superinfection. The Lancet infectious diseases. 2013;13(7):622–628.

